# The C-terminal activating domain promotes Panx1 channel opening

**DOI:** 10.1101/2024.06.13.598903

**Authors:** Erik Henze, Jacqueline J. Ehrlich, Janice L. Robertson, Toshimitsu Kawate

## Abstract

Pannexin 1 (Panx1) constitutes a large pore channel responsible for the release of ATP from apoptotic cells. Strong evidence indicates that caspase-mediated cleavage of the C-terminus promotes the opening of the Panx1 channel by unplugging the pore. However, this simple pore-plugging mechanism alone cannot account for the observation that a Panx1 construct ending before the caspase cleavage site remains closed. Here, we show that a helical region located immediately before the caspase cleavage site, referred to as the "C-terminal activating domain (CAD)," plays a pivotal role in facilitating Panx1 activation. Electrophysiology and mutagenesis studies uncovered that two conserved leucine residues within the CAD plays a pivotal role. Cryo-EM analysis of the construct ending before reaching the CAD demonstrated that the N-terminus extends into an intracellular pocket. In contrast, the construct including the CAD revealed that this domain occupies the intracellular pocket, causing the N-terminus to flip upward within the pore. Analysis of electrostatic free energy landscape in the closed conformation indicated that the intracellular side of the ion permeation pore may be occupied by anions like ATP, creating an electrostatic barrier for anions attempting to permeate the pore. When the N-terminus flips up, it diminishes the positively charged surface, thereby reducing the drive to accumulate anions inside the pore. This dynamic change in the electrostatic landscape likely contributes to the selection of permeant ions. Collectively, these experiments put forth a novel mechanism in which C-terminal cleavage liberates the CAD, causing the repositioning of the N-terminus to promote Panx1 channel opening.

## INTRODUCTION

Pannexin1 (Panx1) forms a large pore membrane channel that releases cellular metabolites and signaling molecules, such as ATP, spermidine, creatine, and lactate (2, 3). Pharmacological intervention and genetic ablation of Panx1 suggest that this channel plays various pathophysiological roles throughout the body (4–6). For example, signaling metabolites released through Panx1 control T cell activation (3), initiate inflammation (7, 8), increase vascular permeability (9), and recruit monocytes for engulfing apoptotic cells (10). Panx1 also contributes to anoxic depolarizations in response to ischemia in the brain (11, 12). This large pore forming channel therefore is an emerging drug target for multiple challenging diseases including cancer and chronic pain (4, 13).

In living cells, Panx1 is activated by a variety of stimuli including membrane stretch/shrinkage (14, 15), voltage (16), increased concentrations of cations (14, 17, 18), post-translational modifications (11, 19), and stimulation of membrane receptors (20–22). Our group recently discovered that lysophospholipids, signaling molecules produced by lipolytic enzymes like phospholipase A2 (PLA2), directly and reversibly activate Panx1 (8). Since PLA2 can be activated downstream of diverse membrane receptors, it is possible that this mode of activation commonly takes place in living cells, although how lysophospholipids activate the channel remains elusive.

In apoptotic cells, it is well-established that the Panx1 channel undergoes direct activation through caspase-mediated removal of the C-terminus (23, 24). This channel-opening mechanism plays a crucial role in releasing cellular metabolites from dying cells, with a particular emphasis on ATP, which serves as a "find-me" signal for recruiting monocytes (10). Cross-linking experiments indicate that the C-terminal region is flexible and the last residue, C426, can reach F54 located in the extracellular side of transmembrane (TM) helix 1 (23). In addition, the substituted cysteine accessibility method uncovered that modifying the C-terminal residues affects Panx1 channel properties (25). Based on these observations, it has been proposed that the C-terminus acts as a pore plug for Panx1, contributing to the closure of the channel under normal conditions.

A number of cryo-EM structures of Panx1 have been determined in the last few years (26). While various solubilization methods (e.g., detergent vs nanodisc) and three distinct species (human, mouse, and frog) were employed to generate cryo-EM maps, the overall conformations were consistent across the datasets (27–31). A notable exception is the N-terminal domain (NTD), which was poorly defined in most datasets. In instances where stronger NTD densities were observed, two distinct conformations were identified: a helix flipped up within the pore (29, 30) and an extended loop flipped down toward the cytoplasm (27, 30). Notably, the flipped-up NTD was associated with Panx1 constructs exhibiting channel activity, while the flipped-down NTD was observed with either inactive constructs or those treated with an inhibitor. These findings, coupled with prior electrophysiological studies (32), provide support for the pivotal role of the NTD in Panx1 channel activity.

Despite the difference in the NTD conformations, the most constricted region of the ion permeation pathway in all available structures is formed by the side chains of the seven W74 residues located in the extracellular domain. The predicted hydration diameter of this constriction is about 9 Å in all structures, which is wide enough for signaling metabolites like ATP to permeate. While this is in line with the notion that the channel with the flipped-up NTD represents an open state, it remains perplexing for Panx1 with the flipped-down NTD, which likely represents a closed channel conformation. One possibility is that the pore-plugging mechanism involving the flexible C-terminus, not visualized in the cryo-EM maps, may contribute to channel closure. However, this mechanism alone cannot entirely explain why a construct lacking the C-terminus is capable of closing the channel (e.g., Panx1Δ365)(27, 33). Another suggested mechanism involves the occlusion of the permeation pathway with lipid molecules (30). While this mechanism has been proposed for other large pore-forming channels (34–36), the process by which lipid molecules enter the pore, predominantly formed with hydrophilic residues, remains unclear. Additionally, the mechanism by which lipid molecules exit the permeation pathway during channel opening requires further investigation.

To better understand the mechanisms underlying the opening and closing of the Panx1 channel, we investigated the role of the C-terminal region important for caspase-mediated activation. Using a combination of truncation, mutagenesis, and electrophysiology techniques, we identified a specific domain within this region that actively facilitates the opening of the Panx1 channel. Additionally, we employed cryo-EM single molecule reconstruction, molecular docking, and continuum electrostatics ion binding free energy calculations to propose a novel mechanism of how the Panx1 channel maintains its closed conformation despite having a wide permeation pathway.

## RESULTS

### The C-terminal activation domain facilitates Panx1 channel activity

The previous cryo-EM structures uncovered that the C-terminal region of Panx1 harbors a well-ordered domain with four internal helices (IH3-6) prior to the caspase cleavage site (Fig. 1A and B). The density between IH6 and the caspase site is relatively weak in currently published maps, although another helix was modeled between I361 and L370 in some structures (29). Whole cell patch clamp recordings from HEK293 cells expressing truncated Panx1 (unless otherwise specified, we used the human orthologue throughout this study) showed that the construct ending at the caspase cleavage site (Panx1Δ379) was substantially more active than wild type, highlighting that removal of the C-terminus promotes Panx1 channel activity (Fig. 1C). This is not specific to voltage stimulus, as lysophospholipid LPC16:0 also facilitated the activity of Panx1Δ379 (Fig. 1D and E). On the other hand, the construct ending immediately after IH6 (Panx1Δ356) failed to show any voltage-dependent channel activity (Fig. 1C), which is consistent with the previous observation (33).

**Figure 1.**
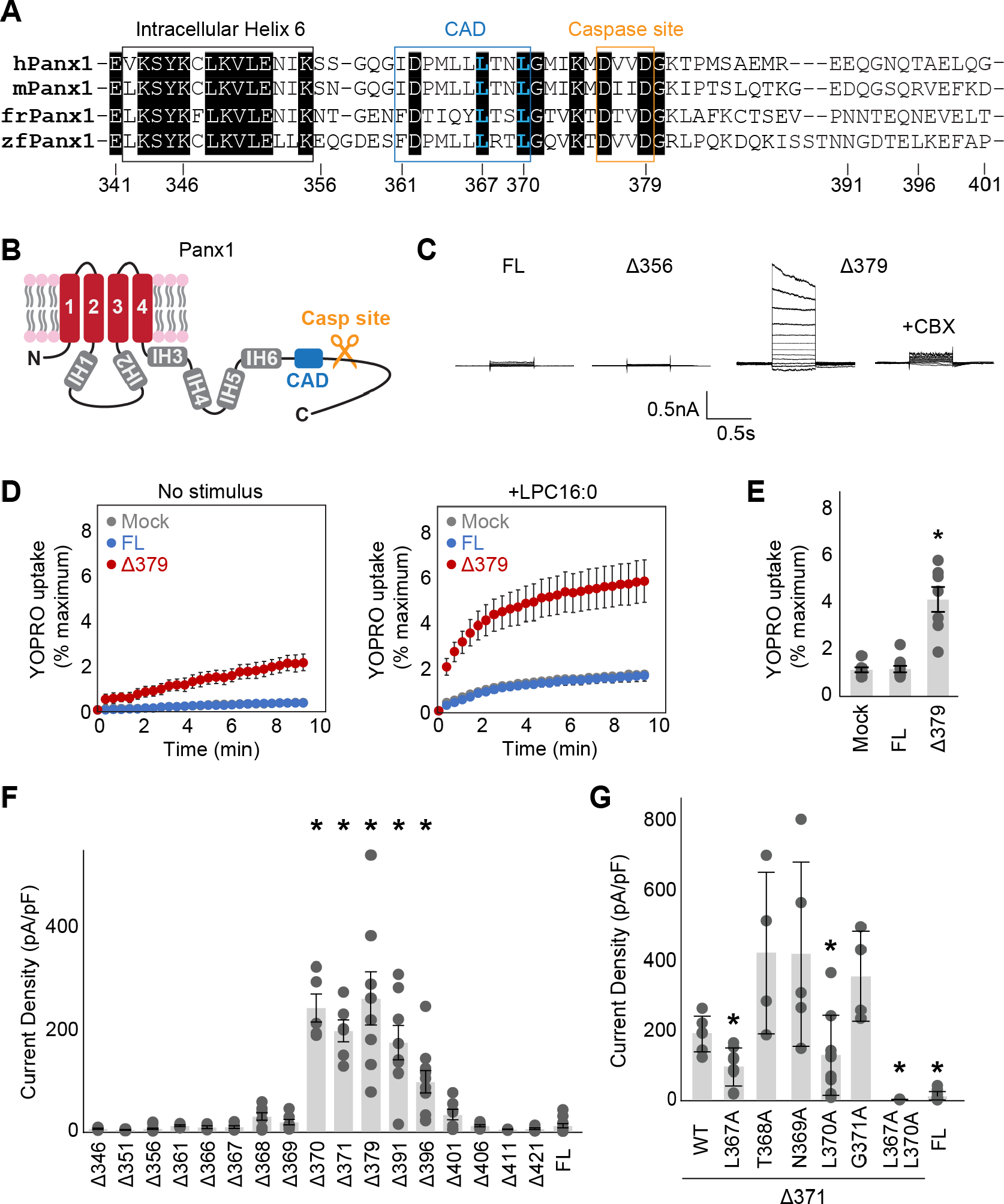
**Systematic truncation uncovers a C-terminal activating domain.** (A) Sequence alignment of the listed Panx1 orthologues around the CAD. (B) A schematic representation of the full length Panx1. IH stands for internal helix. (C) Exemplar whole cell patch clamp recordings of the wild type and C-terminally truncated Panx1 constructs. Cells were clamped at -70 mV and stepped from -110 mV to +110 mV for 0.5 s in 20 mV increments. CBX was applied at 100 μM. (C) and (D) YO-PRO-1 uptake triggered by LPC16:0 (10 μM). Averaged YO-PRO-1 uptake (D) and comparison of the signals at 5 min (E) are shown. N=7-10 and error bars indicate SEM. (F) and (G) Peak whole-cell current density at +110mV obtained from the Panx1 truncation constructs (F) and alanine substitutions of Panx1Δ371 (G). N=4-16 and error bars indicate SEM. Asterisks indicate significance of p<0.05 determined by one-way ANOVA followed by Dunnett’s test comparing the full length Panx1 (FL) to each construct.

We hypothesized that the region spanning from IH6 to the caspase site harbors a domain that facilitates Panx1 channel activity. To identify such a domain, we systematically truncated the C-terminus and tested their activities using whole cell patch clamp recordings. We found that the constructs truncated anywhere before L370 were silent, whereas the ones that end between L370 and T396 showed voltage-activated currents (Fig. 1F). Since the residues between I361 and L370 may form a helix based on the previous cryo-EM maps (29) and circular dichroism experiments (37), we propose that this region forms a structured-domain, dubbed "C-terminal activating domain (CAD)", and mediates Panx1 channel opening. Notably, the constructs ending at G401 or later were also silent, which is consistent with the prevalent idea that the Panx1 C-terminus suppresses its channel activity. To identify the residues in the CAD that are required for facilitating Panx1 channel activity, we systematically examined alanine substitution constructs based on Panx1Δ371 that responds to voltage stimulation. Among those constructs, L367A and L370A diminished their channel activity (Fig. 1G). The double alanine substitutions at these two sites (L367A/L370A) almost completely abolished the currents, supporting that these two residues play pivotal roles in Panx1 facilitation.

We next tested whether the double alanine substitution hinders the opening of the Panx1 channel when activation is triggered by enzymatic C-terminal cleavage. To make proteolysis specific to the Panx1 construct, we substituted the caspase site with the TEV protease site as previously done by Sandilos et al (23). The resulting construct, named Panx1-T (Fig. 2A), exhibited robust voltage-dependent activation only when co-expressed with TEV protease (Fig. 2B and C). In contrast, Panx1-T L367A/L370A did not respond to voltage stimulation even in the presence of TEV protease. We observed the same trends with inside-out patch recordings where C-terminal cleavage was initiated with perfusing a recombinant TEV protease (Fig. 2D and E). These experiments confirm that L367 and L370 in the CAD play essential roles in Panx1 activation triggered by enzymatic C-terminal cleavage.

**Figure 2.**
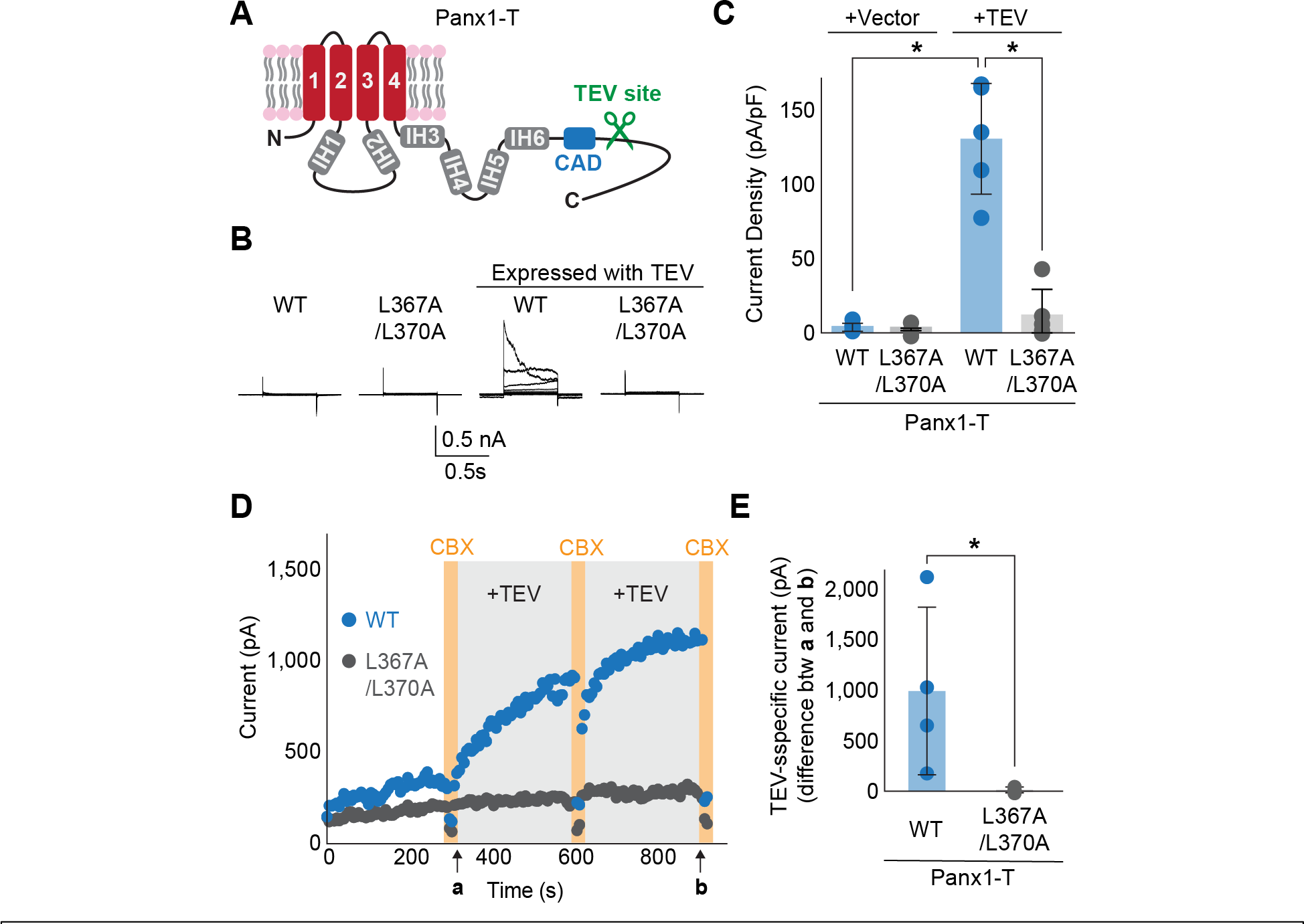
**Enzyme-mediated C-terminal cleavage exposes the CAD.** (A) A schematic representation of the full length Panx1 harboring a TEV protease cleavage site instead of the caspase recognition site (Panx1-T). (B) and (C) Exemplar whole cell currents (B) and peak current density at +110 mV (C) of Panx1-T co-expressed in HEK293 cells with TEV protease. Cells were clamped at -70 mV and stepped from -110 mV to +110 mV for 0.5 s in 20 mV increments. N=4. Asterisks indicate significance of p<0.05 determined by student T-test. (D) Inside-out patch clamp recordings with or without recombinant TEV protease (0.1mg/mL). Voltage ramps from 130 mV to +80 mV over 0.5 s were applied every 6 s. Representative peak currents at +80 mV are plotted for Panx1-T (blue) and Panx1-T L367A/L370A (gray). CBX (100 μM) were applied at time points indicated by orange bars. (E) TEV specific currents defined as the peak current differences between the indicated time points **a** and **b**. N=4. Asterisk indicates a p<0.05 obtained from an unpaired student T-test.

### Freed CAD promotes NTD flip

Considering the positive action of the CAD on Panx1 activation, we hypothesized that the exposure of this domain following caspase cleavage induces a conformational change essential for channel opening. To visualize the CAD-driven conformational change, we compared the cryo-EM structures of two Panx1 constructs, one terminating before and the other after this domain. We used the frog orthologue for this structural analysis due to its superior expression and stability (27). To retain both the N- and C-termini, we introduced a Strep affinity tag in the flexible intracellular loop between IH1 and IH2. The functionality of the resulting constructs, termed frPanx1-ΔC and frPanx1-ΔC+CAD (Fig. 3A), was assessed via YO-PRO-1 uptake assay. Upon LPC16:0 treatment, frPanx1-ΔC+CAD exhibited robust dye uptake, which was effectively inhibited by its antagonist spironolactone (Fig. 3B). Conversely, neither vector-transfected cells nor frPanx1-ΔC displayed noticeable dye uptake. These findings confirm that frPanx1-ΔC remains in its closed conformation, while frPanx1-ΔC+CAD adopts an open or facilitated conformation, poised to respond to its stimulus.

**Figure 3.**
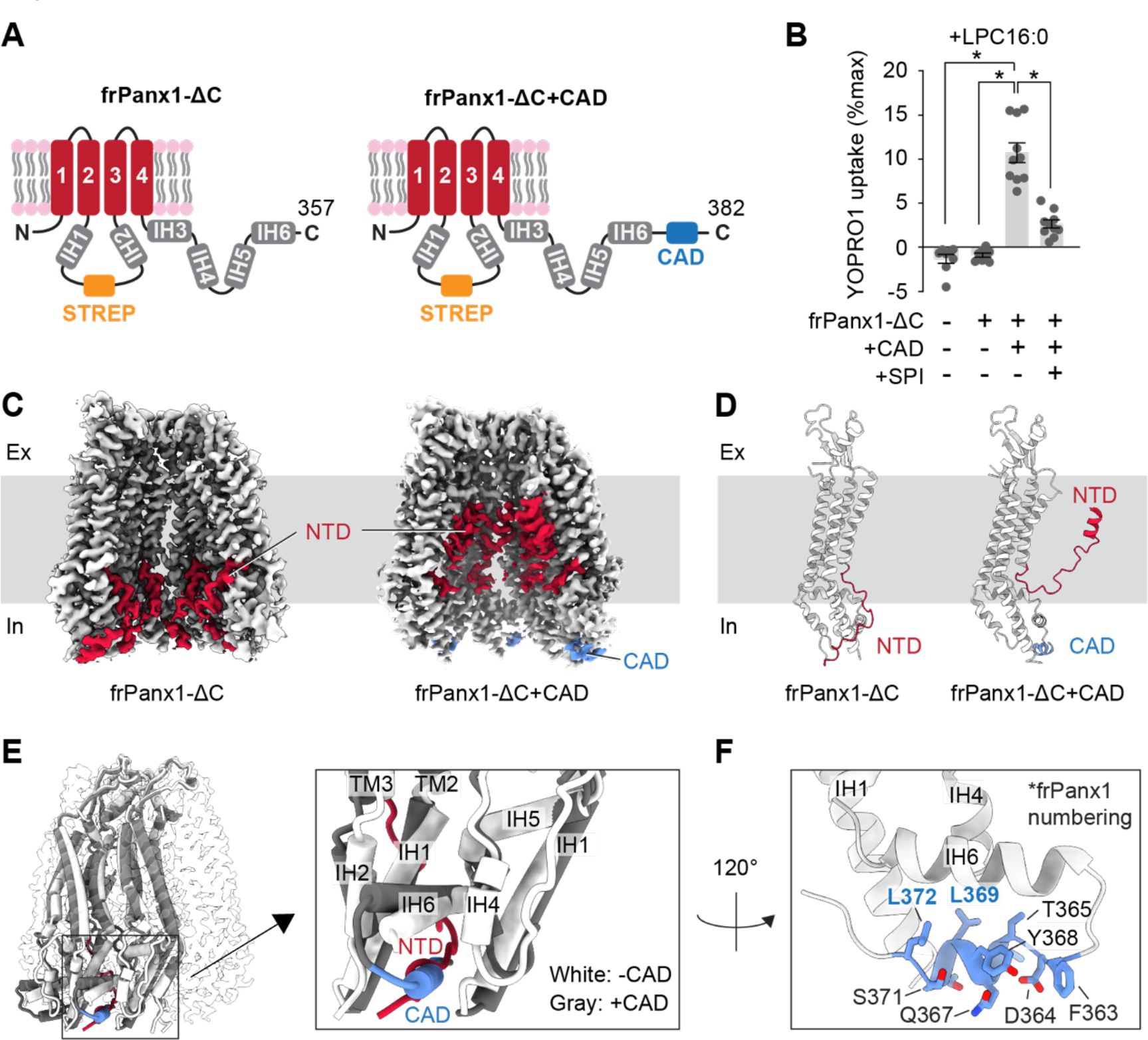
**CAD promotes the NTD to flip.** (A) Schematic representations of frPanx1-ΔC (left) and frPanx1-ΔC+CAD (right). Residues between 171 and 179 were replaced with StrepII tag. (B) YO-PRO- 1 uptake triggered by LPC16:0 (10 μM). YO-PRO-1 uptake from frPanx1 expressing Sf9 cells at 5 min are shown. N=10 and error bars indicate SEM. Asterisks indicate significance of p<0.05 determined by one-way ANOVA followed by Dunnett’s test comparing to frPanx1-ΔC+CAD. SPI; spironolactone (50 μM). (C) and (D) Cryo-EM structures of the frPanx1 constructs. Cryo-EM maps (C) and cartoon representation of a protomeric model of frPanx1-ΔC (left) and frPanx1-ΔC+CAD (right). EM densities for only four subunits are shown for clarity. (E) Superposition of frPanx1-ΔC (while) with frPanx1- ΔC+CAD (gray) highlighting that the NTD (red) and the CAD (blue) occupies a common binding pocket. (F) Close-up view of the frPanx1-ΔC+CAD model around the CBP.

The cryo-EM structure of frPanx1-ΔC revealed a flipped-down N-terminal domain (NTD) (Fig. 3C and D; Fig. S1, 3 and 4; table S1), consistent with previously observed structures in its closed state (27, 30). In contrast, frPanx1-ΔC+CAD exhibited a flipped-up NTD with helices aligned in the pore (Fig. 3C and D; Fig. S2- 4; table S1). Importantly, the CAD occupies a pocket between neighboring subunits, termed the "CAD-binding pocket (CBP)," where the NTD resides in its closed conformation as observed in the frPanx1-ΔC structure (Fig. 3E). While the EM density of the CAD was too weak to confidently assign side chain residues, the two critical residues in this domain, L367 and L370 (L369 and L372 in frPanx1, respectively), face the hydrophobic surface of the CBP (Fig. 3F). This finding suggests that the liberated CAD following caspase cleavage replaces the NTD situated within the CBP, facilitating the flipping of this domain into the pore.

If the CAD operates by competing with the NTD, we reasoned that this domain does not necessarily need to be situated at its natural position, as long as it is placed near the CBP. To test this idea, we introduced the CAD into the flexible intracellular loop before IH2 in the Panx1Δ356 background, which on its own does not respond to voltage stimulus (Panx1Δ356+CAD-IL1; Fig. 4A). Whole-cell patch clamp experiments revealed that incorporating the CAD into the intracellular loop facilitated voltage-dependent channel opening (Fig. 4B and E). Notably, the L367A/L370A double mutant failed to activate, indicating that the ectopically positioned CAD retains its functionality. Interestingly, when the CAD was inserted near IH1 in the intracellular loop (Panx1Δ356+CAD-IL2; Fig. 4C), the channel did not respond to voltage (Fig. 4D and E). Considering that IH1 extends deeper into the cytoplasm than IH2, it is likely that the CAD could not reach the CBP. Indeed, when a flexible seven-amino acid linker (GSGSGSG) was inserted between IH1 and the CAD, we observed voltage-stimulated channel activity dependent on L367/L370 (Fig. 4D and E). We also investigated whether full length Panx1, which does not respond to voltage stimulation (Fig. 1C), could be activated by an ectopically inserted CAD. Whole-cell recordings revealed that the CAD inserted at intracellular loop position 1 (Panx1- FL+CAD-IL1) and at the very end of the C-terminus (Panx1-FL+CAD-CT) were activated by voltage (Fig. 4F-J). These findings collectively support that the CAD can facilitate Panx1 activation even when located in non-native positions.

**Figure 4.**
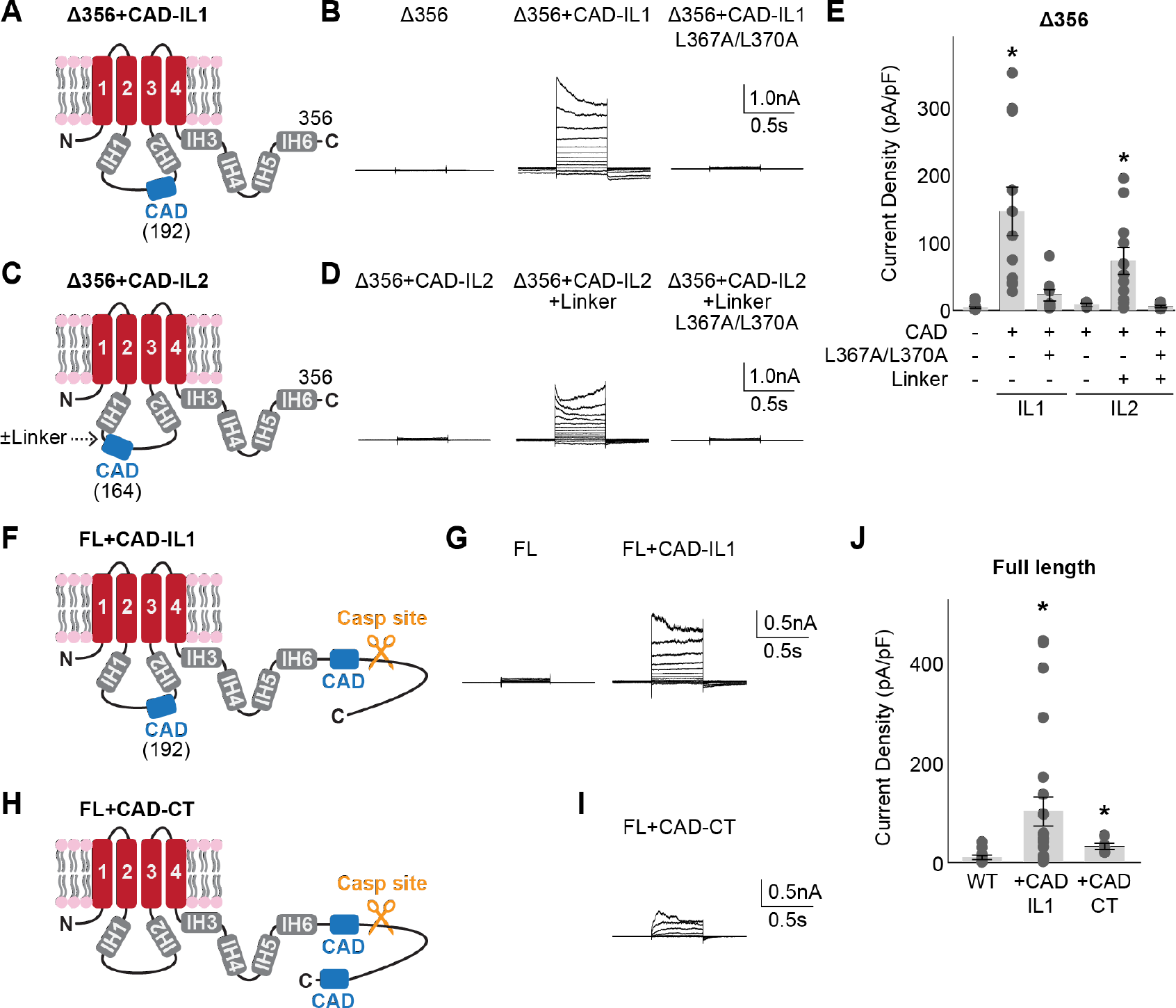
**CAD can facilitate Panx1 activation from non-native positions.** Schematic representations ((A), (C), (F), and (H)), representative whole cell recordings ((B), (D), (G), (I)), and peak current density at +110 mV ((E) and (J)) are shown for each tested construct. HEK293 cells were clamped at -70 mV and stepped from -110 mV to +110 mV for 0.5 s in 20 mV increments. N=5-24 and error bars indicate SEM. Asterisks indicate significance of p<0.05 determined by one-way ANOVA followed by Dunnett’s test comparing the Panx1Δ356 (E) or Panx1 wild type (J) to each construct.

### The effects of certain posttranslational modifications depend on CAD activity

Previous studies have suggested that Panx1 channel activity is modulated by posttranslational modifications (PTMs). For example, S-nitrosylation at C40 and C347 or acetylation at K140 downregulates its activity, while phosphorylation at Y150, Y199, or Y309 enhances its activity (12, 19, 38–41). Indeed, blocking inhibitory-PTMs or mimicking the facilitatory-phosphorylation through mutagenesis renders Panx1 responsive to voltage stimulation (Fig. 5A and B). Notably, these PTM sites, except for C40, are clustered near the CBP (Fig. 5C). To assess whether the effects of these PTMs depend on CAD function, we introduced L367A/L370A double mutations in the Panx1 PTM point mutants. Voltage-stimulated channel activity of the three point mutants blocking the PTMs (i.e., C40S, K140R, and C347S) was significantly reduced by L367A/L370A mutations, suggesting that the Panx1 facilitation by these mutations depend on CAD activity (Fig. 5D). Likewise, one of the three phosphomimetics, Y199D, lost its activity with the L367A/L370A mutations. In contrast, the other two phosphomimetics, Y150D and Y309D, remained active even in the presence of the L367A/L370A mutations, indicating that phosphorylation at these residues potentiates Panx1 channel activity independent of the CAD. In all cases, truncation of the first 20 residues in the NTD completely abolished its channel activity (Fig. 5D). This effect is not due to impaired surface expression, as indicated by the comparable or even higher plasma membrane protein levels of each mutant with the NTD truncation (Fig. S5). Together, these experiments suggest that previously identified PTMs influence the movement of the NTD, either through direct action or via CAD function, to regulate Panx1 activity.

**Figure 5.**
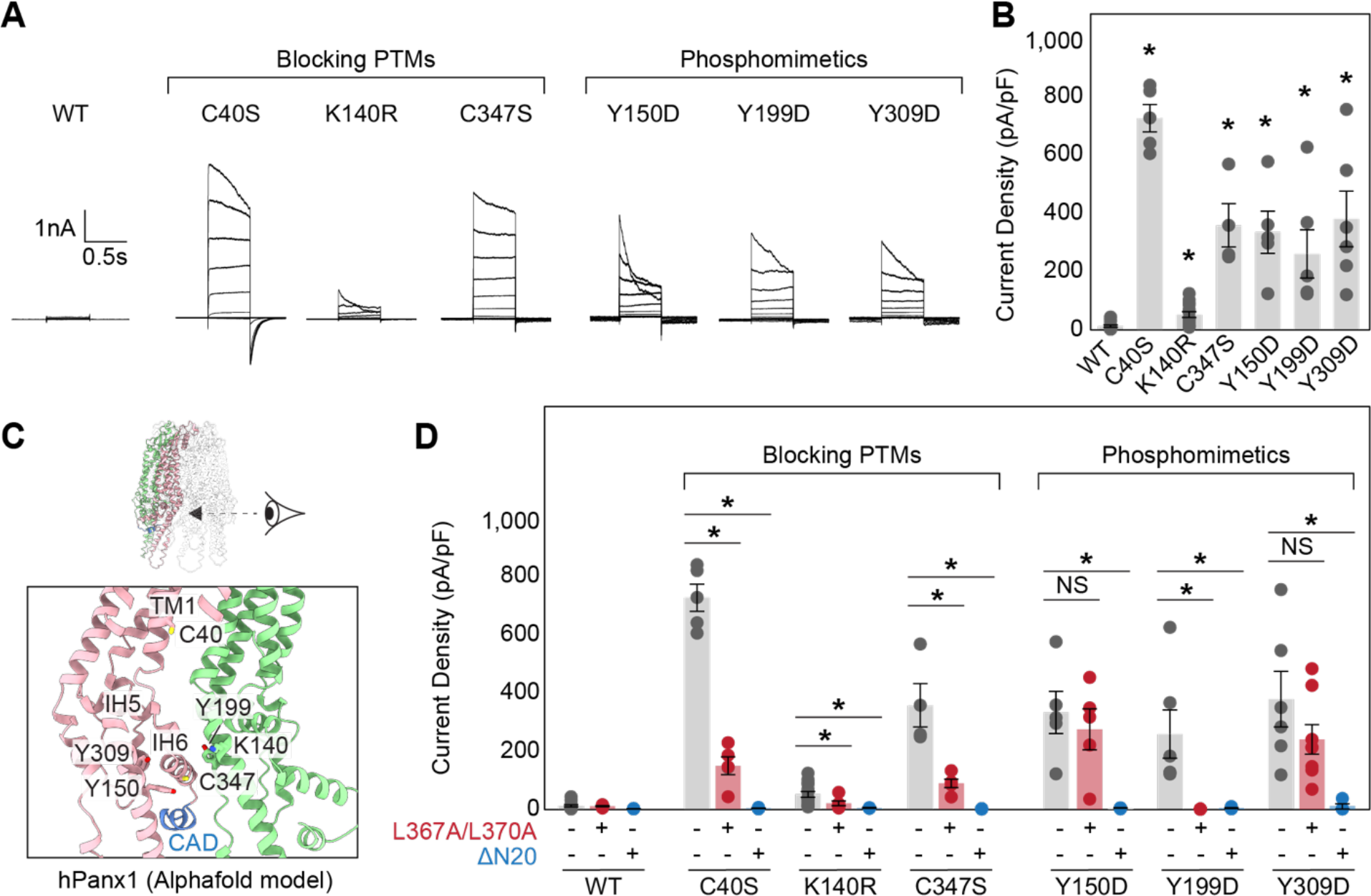
**Point mutants blocking or mimicking PTMs potentiate Panx1 activity through CAD and NTD.** Exemplar whole cell patch clamp recordings (A) and peak current density at +110 mV (B) of the wild type and point mutants. Cells were clamped at -70 mV and stepped from -110 mV to +110 mV for 0.5 s in 20 mV increments. N=4-8 and error bars indicate SEM. Asterisks indicate significance of p<0.05 determined by one-way ANOVA followed by Dunnett’s test comparing WT to each construct. (C) Cartoon representation of the human Panx1 structure around the CBP. The AlphaFold model of human Panx1 was used due to missing side chains in the currently available cryo-EM structures. Only two subunits are shown for clarity. (D) Comparison of the peak current densities for wild type and each point mutant with or without L367A/L370A mutations or N-terminal truncation (2–20). N=4-8 and error bars indicate SEM. Asterisks indicate significance of p<0.05 determined by one-way ANOVA followed by Dunnett’s test comparing the parent construct with the L367A/L370A mutations and N-terminal truncation for each point mutation.

### Anion occupancy in the permeation pathway contributes to the channel closure

How does the NTD flip promote Panx1 channel opening? Given that the narrowest portion of the permeation pathway in the closed frPanx1-ΔC channel (NTD flipped down) is approximately 9 Å in diameter, a steric barrier mechanism alone cannot explain how this channel prevents ion permeation. Therefore, we compared the electrostatic surface potentials between the two conformations with the NTD flipped down (frPanx1-ΔC) and flipped up (frPanx1-ΔC+CAD). We observed that the extracellular region near the narrowest point of the permeation pathway (around W74) in the NTD flipped-down conformation carries more negative charges. This suggests the potential role of this region as an electrostatic barrier for anion permeation (Fig. 6A; left). Notably, in the NTD flipped-up conformation, this region transitions to a neutral to slightly positive charge (Fig. 6A; right). Additionally, we found that the middle section of the permeation pathway exhibits a strong basic potential in the flipped-down conformation, while it becomes slightly acidic in the flipped-up conformation (Fig. 6A; left).

**Figure 6.**
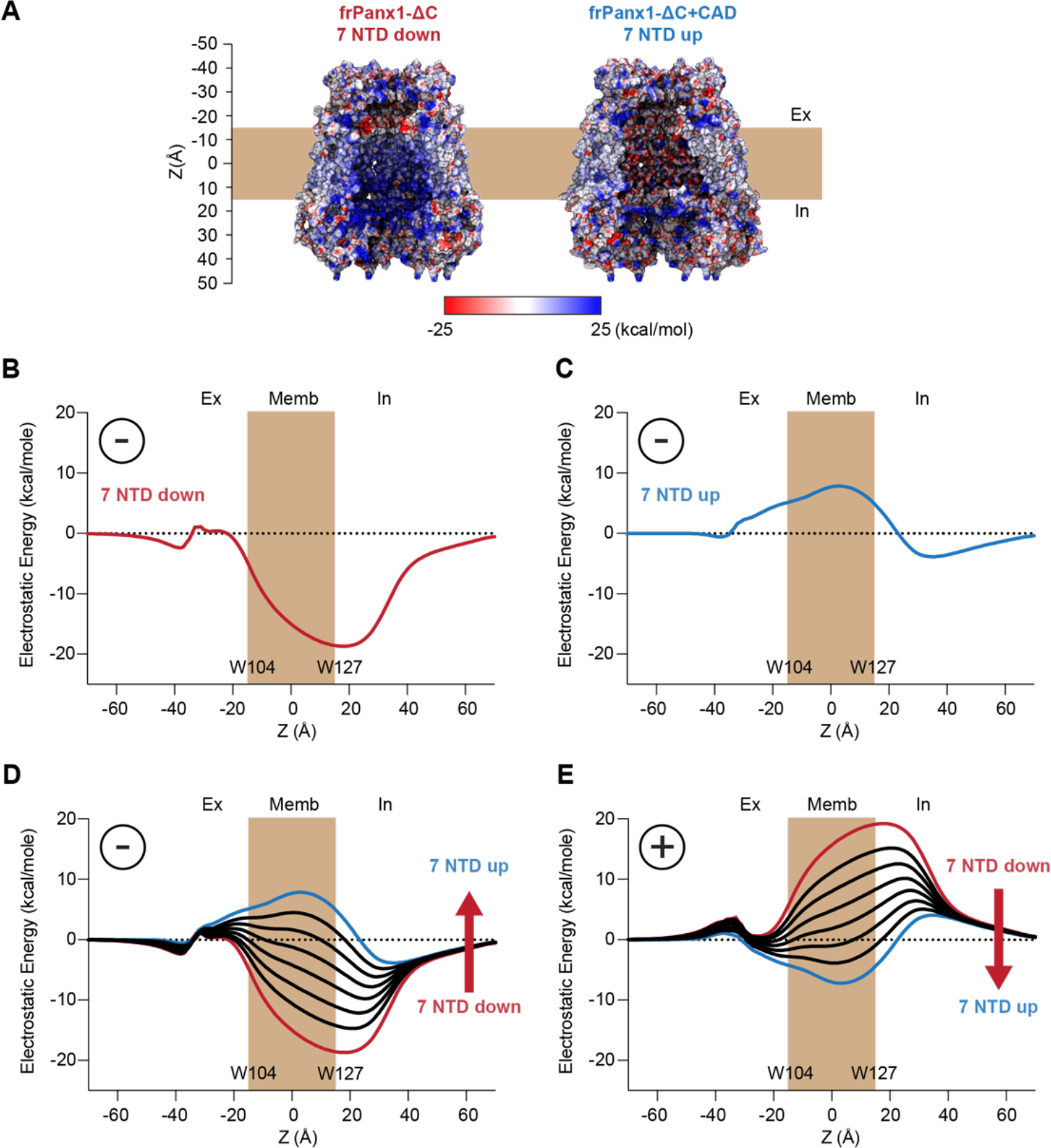
**Electrostatic energy landscape controls ion permeation through the Panx1 channel.** (A) Electrostatic surface potential of a coronal section of the frPanx1-ΔC (left) and frPanx1-ΔC+CAD (right) ion permeation pathway. Electrostatic surface potential was calculated using the PBEQ module in CHARMM (1) and presented in the range between -25 kcal/mol (red: acidic) and +25 kcal/mol (blue: basic). Electrostatic contribution for an anion binding free energy (Etotal) along the permeation pathway (z=0 in the middle of predicted transmembrane) for the frPanx1-ΔC (B), frPanx1-ΔC+CAD (C), and hybrid models harboring 0-7 NTDs (D). (E) Electrostatic contribution for a cation binding free energy along the permeation pathway for the hybrid models. All calculations are carried out using the PBEQ module in CHARMM to solve the Poisson-Boltzmann solution of the system.

To further investigate how changes in the pore could impact ion permeation, we calculated the change in the electrostatic free energy associated with transferring an anion from the bulk into the central pore axis as a function of the Z position. For reference, the structures are aligned so that W104 is located at one side of the membrane (Z ≈ -17 Å) and W127 at the other side (Z ≈ 17 Å). This calculation includes the contribution from the ion in the protein static field, described by the surface potential maps in Fig. 6A, but also includes the ion reaction field, a repulsive term dependent on the protein and membrane dielectric boundary (Fig. S6A and B). Since the channel is wide open in both conformations, the reaction field contribution is small except around the constriction position of W74 (Fig. S6C and D). In the NTD flipped down structure (frPanx1-ΔC), the pore is overwhelmingly favorable for anions, whereas the NTD flipped up structure (frPanx1- ΔC+CAD) reveals a slight barrier (Fig. 6B and C). While this appears to contradict the postulated closed and open characteristics, it is important to remember that this analysis provides a scan of the electrostatic character of the pore but lacks the context that is relevant to permeation. Thus, we also investigated how modulation of both the protein structure and ion occupancy alter these electrostatic profiles.

The NTD flipped-up structure is predicted to form the conductive pore, but our results indicate a slight barrier for anions. We speculate that under active conditions, there are fractional occupancies of the NTD in their sites rather than all 7 NTD being flipped up. To test this idea, we carried out the same calculations as a function of the number of NTD in the up position by building hybrid structural models. Flipping just 1-2 NTDs down reduces the barrier close enough to zero supporting permeation (Fig. 6D). These profiles also correspond to the 7 NTD flipped-up electrostatic profile of the human Panx1 (Fig. S7A and B). In addition, it stands that cation permeation may be favorable in the NTDs up configuration (Fig. 6E). Thus, considering that NTD occupancy may be variable, the electrostatic profiles with at least 3 NTDs bound, would support ion permeation of some form. Thus, this structure is in line with a conductive pore that is generally non-selective or demonstrates differential selectivity depending on the NTD occupancy.

On the other hand, the NTD flipped down structure is postulated to be a closed channel, and it contains a deep electrostatic well around Z = 20 Å (Fig. 6B). Such an environment likely promotes the accumulation of anions, potentially multivalent ones such as ATP. To assess how ATP molecules in the deep well would influence ion permeation, we predicated a possible binding position using molecular docking and calculated the electrostatic free energy for anion permeation as a function of the number of ATP molecules docked (Fig. 7A and B). As anticipated, the depth of the energy well decreased with the increasing number of ATP molecules placed (Fig. 7C). Remarkably, we noticed that the negative charges of ATP molecules contributed to the formation of an electrostatic energy barrier on the extracellular side, near residue W74. This barrier became prominent with just two ATP molecules within the deep well and intensified with three molecules. Considering ATP’s abundance in the cytoplasm at millimolar concentrations, it is plausible that this molecule binds to the deep energy well in the NTD flipped-down conformation, thereby impeding anion permeation. This channel closure mechanism is not specific to the frog species, as we observed the same phenomenon with the human Panx1 structures representing the NTD flipped-down (PDB: 7F8N) and flipped-up (PDB: 7F8J) conformations (Fig. S7C).

**Figure 7.**
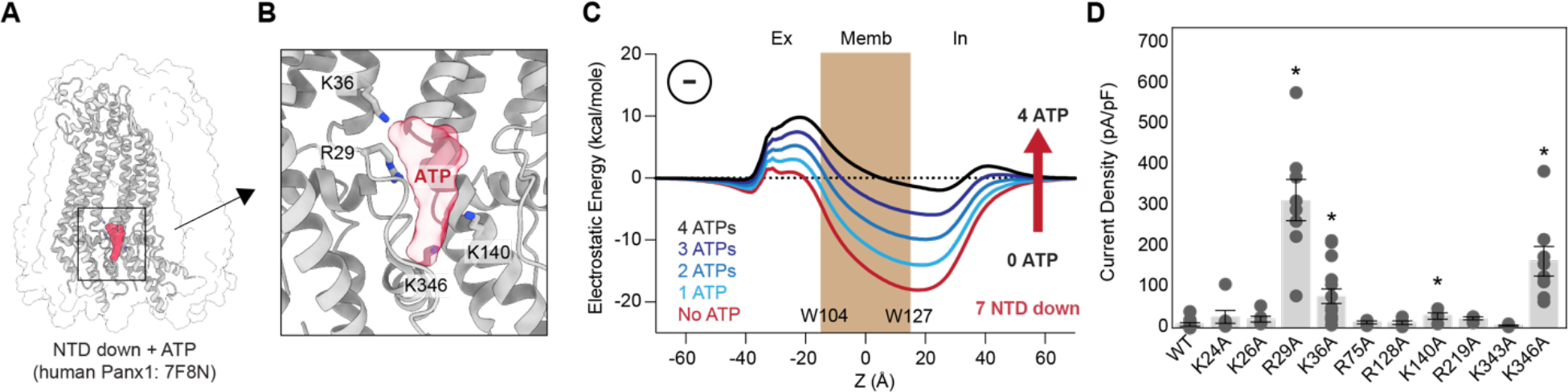
**Docked anions in the permeation pathway contribute to the channel closure.** (A) Electrostatic free energy calculation of frPanx1-ΔC (red) in the presence of 1-4 ATP molecules in the deep energy well. (B) A docked ATP molecule on the NTD-flipped down human Panx1 (PDB: 7F8N). (C) Close-up view of the putative ATP binding pocket. Positive residues surrounding the docked ATP are indicated. (D) Peak current density at +110 mV of the wild type and alanine substitutions expressed in HEK293 cells. Cells were clamped at -70 mV and stepped from -110 mV to +110 mV for 0.5 s with 20 mV increments. N=4-15 and error bars indicate SEM. Asterisks indicate significance of p<0.05 determined by one-way ANOVA followed by Dunnett’s test comparing WT to each construct.

To further explore the impact of the deep energy well on ion permeation, we investigated the Panx1 channel activity of point mutants targeting the positively charged residues surrounding the well. Among the ten alanine substitution mutants in this region, R29A, K36A, and K346A notably enhanced the voltage-dependent channel activity (Fig. 7D). This supports that the positive charge in this region contributes to the channel closure. Although cytoplasmic ATP is likely absent in our whole-cell recording conditions, the high concentrations of Cl^-^ and HEPES ions present would serve a similar function in generating the extracellular electrostatic barrier. These findings collectively suggest that the deep energy well in the NTD-flipped down conformation hinders anion permeation by attracting anions, thereby acting as an electrostatic energy barrier. Flipping up the NTP disrupts anion binding in this area, facilitating anion permeation. Our current Panx1 activation and ion selection models are summarized in Figure 8.

**Figure 8.**
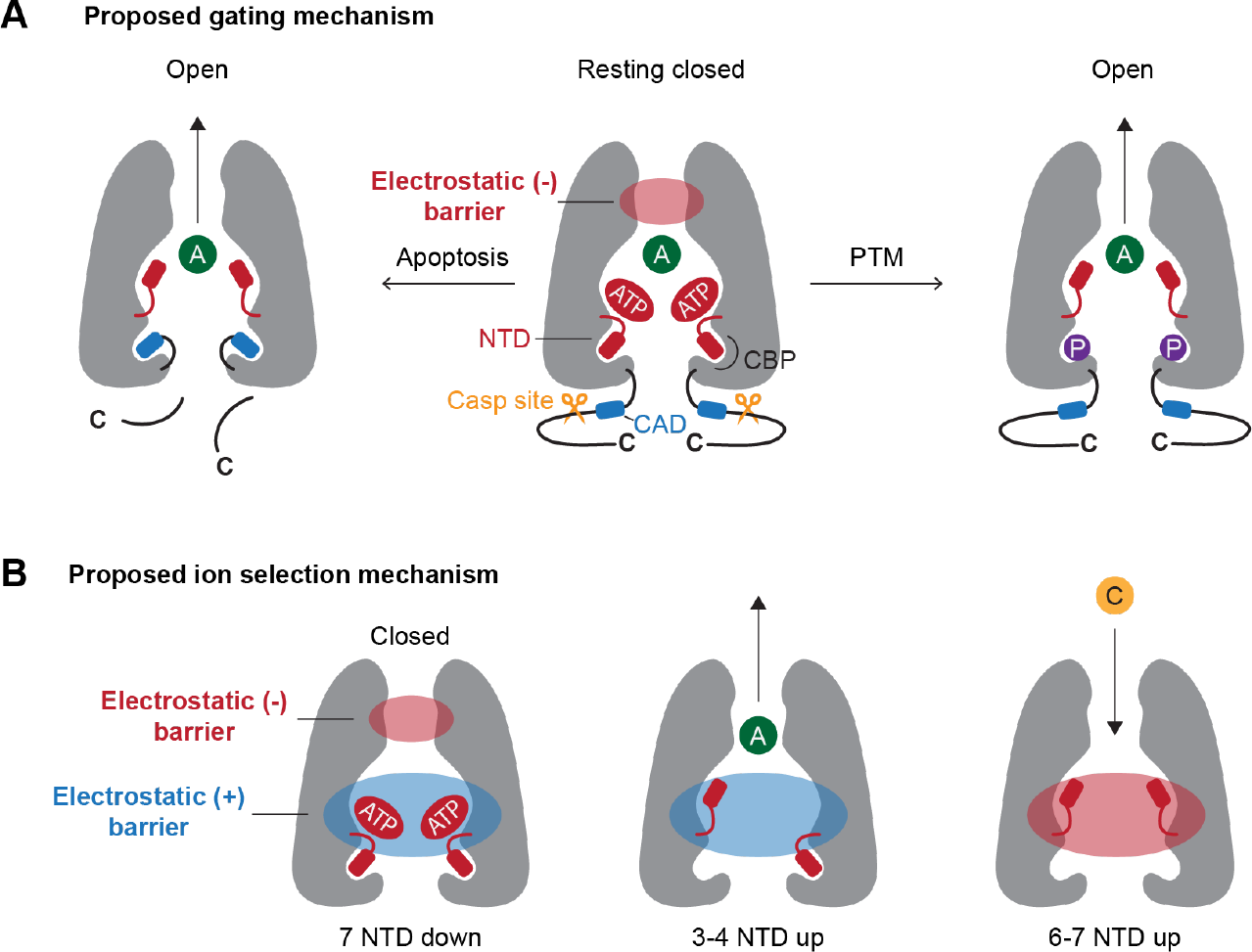
**Current models for Panx1 gating and ion selectivity.** (A) At rest, the NTD extends into the cytoplasm, which contribute to generate highly basic surface attracting anions such as ATP. The high occupancy of anions contribute to create an electrostatic energy barrier at the extracellular side of the permeation pathway, thereby limiting the permeation of anions (middle). In apoptotic cells, caspase cleavage liberates the CAD, triggering the flipping of the NTD (right). Certain PTMs (e.g. phosphorylation at Y150D) promote the NTD flip to remove the electrostatic energy barrier (left). (B) Both negative and positive electrostatic barriers limit the permeation of ions in the closed state (left). When 3-4 NTDs flip up, the negative electrostatic barrier disappears, facilitating anion permeation (middle).

## DISCUSSION

In this study, we have demonstrated that cleaving the Panx1 C-terminus by caspase exposes a helical domain named the "C-terminal activating domain (CAD)", which facilitates Panx1 channel opening. Mutating two leucine residues within the CAD, L367 and L370, nearly abolishes Panx1 channel activity induced by C-terminal cleavage, underscoring the critical roles of these hydrophobic residues in CAD function. The Panx1 channel truncated upstream of the CAD remains closed, even in the presence of an activation stimulus. In this state, the NTD is positioned downward into the cytoplasm, while the addition of CAD competes for this binding pocket and flips up this domain within the permeation pathway. Based on our electrostatic free energy calculations, we propose that the NTD’s downward conformation attracts anions around the cytoplasmic side of the permeation pathway, which generates an electrostatic barrier on the extracellular side. NTD flipping, triggered either by the liberated CAD following caspase cleavage or by posttranslational modifications, disrupts the anion binding site, thereby reducing the electrostatic barrier for anion permeation (Fig. 8).

In the resting closed conformation, it has been demonstrated that the intact Panx1 C-terminus plugs the pore (23, 29). According to our current model, the C-terminus likely plays an additional role in channel closure by obstructing the access of the CAD to its binding pocket (CBP). Given that none of the currently available cryo-EM structures of full-length Panx1 show conspicuous C-terminal density beyond the caspase cleavage site, it is plausible that the mechanism by which this region plugs the pore or sequesters the CAD may involve flexible structures. Indeed, a previous study by Dourado et al. demonstrated that the function of the C-terminal region does not require a specific sequence (33).

Although the mechanism by which the C-terminus sequesters the CAD remains unclear, it is conceivable that this mechanism is linked to pore plugging. When C40 is mutated to serine to prevent post-translational S-nitrosylation, the full-length Panx1 becomes responsive to voltage stimulation (Fig. 5C and D). One possible explanation for this gain-of-function effect is that C40S facilitates the flip of the NTD by attracting this domain to its vicinity. However, this mutant loses its ability to enhance channel activity when L367 and L370 within the CAD are mutated to alanines. Given that C40 is situated in the middle part of the ion permeation pathway away from the CBP, it is unlikely that post-translational modification of this residue directly influences CAD binding. Instead, we speculate that the distal C-terminus, which normally interacts with the S-nitrosylated C40 to plug the pore (38), becomes more relaxed due to the serine substitution, thereby facilitating CAD binding to the CBP.

Previously determined cryo-EM structures of full-length human Panx1 have displayed the NTD-flipped up conformation similar to the frPanx1-ΔC+CAD structure obtained in this study (29, 30). While this observation suggests that the full-length Panx1 channel is in an active conformation, we propose that the channel activity is cell type dependent. In HEK293 cells, both our research and others’ findings have shown that full-length Panx1 does not respond to activation stimuli (42). However, in other cell types such as tsA201 or HEK293S GnTI^-^ cells, this construct exhibits voltage- or lysophospholipid-dependent channel activity (8, 29). Although the underlying mechanism remains elusive, we speculate that different post-translational modifications (PTMs) may either inhibit or promote Panx1 channel activity. Indeed, our current study demonstrated that point mutants blocking or mimicking PTMs can render full-length human Panx1 responsive to activation stimuli, even in HEK293 cells (Fig. 5).

The cryo-EM structures determined in this study agree with previous observations of closed and opened Panx1 conformations. Although we took advantage of the superior expression and solvability of the frog Panx1 ortholog, the overall assembly of frog and human Panx1 are nearly identical. Regions of flexibility in both orthologs are also consistent. However, we noticed that the NTD helix in the frPanx-ΔC+CAD structure makes contacts with the transmembrane helices (TMHs) of the neighboring subunit rather than on its own. This connection contrasts the human Panx1 structure (7F8J) whose N-termini make contacts with the TMHs of its own subunit (30). Given the weak cryo-EM density of the connection between the first TMH and the NTD, we isolated each protomer with 3D masks of each N-terminal connection to better visualize the density in this region. In these isolated Panx1 particles, there was a connection when the mask encompassed the neighboring TMH, and virtually no connection when it covered the TMH of the same subunit (Fig. S4). Although we cannot exclude the possibility that the connections vary among different species, it is more likely that the NTDs are able to interact with the TMHs on either subunit, and they could swap as the channel cycles between open and closed conformations.

Our electrostatic free energy calculations suggest that anions accumulated within the deep energy well of Panx1 contribute to the formation of an electrostatic barrier in the extracellular domain. This resembles to the mechanism in which the ZntB transporter may preferentially select cations over anions for its substrate (43). In this transporter, positively charged residues in the vestibule create an electrostatic barrier that impedes cations from accessing the cytoplasmic entrance of the pore. Conversely, when multiple Cl^-^ ions bind to the vestibule, the electrostatic barrier dissipates, and the entire vestibule becomes attractive for cations. It is conceivable that the deep energy well in Panx1 is typically occupied by anions in the resting closed channel with the NTD flipped down. Considering that most, if not all, Panx1 activity assays have been conducted in the presence of a high concentration of anions, our current research provides an explanation for why a large pore channel with a 9 Å opening can prevent permeant ions from passing through. Nevertheless, it is likely that other mechanisms for Panx1 channel closure exist, as uncharged molecules such as glucose do not seem to permeate the channel (15).

In conclusion, our findings demonstrate that ion permeation in the Panx1 channel is regulated by the electrostatic landscape. We propose that the movement of both the CTD and the NTD significantly influences Panx1 channel gating by modulating these electrostatic properties within the permeation pathway. We anticipate that future studies will elucidate the mechanisms by which Panx1 activators, including voltage and lysophospholipids, trigger channel opening.

## MATERIALS AND METHODS

### Expression constructs

DNA constructs encoding the amino acid sequences of the human Panx1 (hPanx1: gene ID: 24145) and frog Panx1 (frPanx1: 100170473) were synthesized (GenScript) and subcloned into appropriate vectors using a standard molecular cloning techniques. We used pIE2 or pIM2 vectors for transient expression in HEK293 cells and pFB-NT vector for insect cell expression. The human Panx1 CAD sequence (residues 361-370) was inserted at the designated position by PCR. A C-terminal FLAG tag for pannexin western blotting and internal Strep II tag replacing residues 177- 184 for frPanx1-ΔC purification were introduced by PCR. Point mutants were generated by site-directed mutagenesis. A plasmid containing maltose binding protein (MBP) fused with an N-terminal His x6 tagged TEV protease (S219V) was gifted from Josh Chappie. The coding region of the fusion protein was subcloned into pNNG-BC10 vector for bacterial expression using PCR. All constructs were verified by sequencing.

### Cell culture

HEK293 cells (ATCC: CRL-1573) were cultured in DMEM supplemented with 10% FBS at 37°C with 8% CO2 in a humidified incubator. The mycoplasma contamination test was confirmed to be negative at ATCC and therefore were not further authenticated. Sf9 cells (ThermoFisher Scientific: 11496015) were maintained in Sf-900 III SFM (ThermoFisher Scientific) and HighFive cells (ThermoFisher Scientific: B85502) were maintained in ESF 921 (Expression Systems) at 27 °C with a shaking speed of 125 rpm.

### Patch-clamp recordings

HEK293 cells (passage number < 40) were plated at low density onto 12-mm glass coverslips in a six-well plate (Greiner). Cells were transfected after 24 h with 800 ng plasmid DNA using FuGENE6 (Promega) according to the manufacturer’s instructions. Recordings were obtained 40- 60 hours after transfection. Borosilicate glass pipettes (Harvard Apparatus) were pulled and heat polished to a final resistance of 2-4 MΩ and backfilled with (in mM) 147 NaCl, 10 EGTA, and 10 HEPES (adjusted to pH 7.0 with NaOH). Whole cell patches were obtained in an external buffer containing (in mM) 147 NaCl, 2 KCl, 2 CaCl2, 1 MgCl2, 13 glucose, and 10 HEPES (adjusted to pH 7.3 with NaOH). For inside-out recordings, pipettes contained the bath solution. Inside-out patch perfusion buffer contained (in mM) 155 NaCl, 10 HEPES and 0.1 EGTA adjusted to pH 7.0. Inside-out patches were obtained by transient air exposure of patches pulled from Panx1- expressing cells. To measure the effect of TEV protease on Panx1-mediated currents, a voltage ramp protocol (-130mV to +80mV over 0.5s) was applied every 6 seconds. TEV protease (100 μg/mL) was perfused onto the inside-out patch. Every 5 minutes, 100 μM CBX was applied for 2 voltage ramps. A rapid solution exchange system (RSC-200; Bio-Logic) was used for recordings in which patches were perfused with drugs. Currents were recorded using an Axopatch 200B patch-clamp amplifier (Axon Instruments), filtered at 2 kHz (Frequency Devices), digitized with a Digidata 1440A (Axon Instruments) with a sampling frequency of 10 kHz, and analyzed using the pCLAMP 10.5 software (Axon Instruments).

### YOPRO Uptake Assay

HEK293 cells were cultured on 96-well poly-D-lysine coated black-walled plates (Corning: 356640). At 90% confluency, cells were transfected with 200 ng Panx1 DNA using jetPRIME (Polyplus) according to the manufacturer’s instructions. For frPanx1 constructs, Sf9 insect cells were infected with 5% P2 virus for 2 days and plated on 96-well poly-D-lysine coated black-walled plates prior to the assay. After 20 hours incubation growth media was replaced with warmed Assay buffer including (in mM) 147 NaCl, 2 KCl, 2 CaCl2, 1 MgCl2, 13 glucose,10 HEPES (adjusted to pH 7.3 with NaOH), and 5 μM YO-PRO-1. When antagonists were used in experiments, Assay buffer also included 100 μM CBX or 50 μM spironolactone. To stimulate Panx1, LPC16:0 (10 μM) was added after 5 min equilibration. Pannexin activity was monitored by measuring fluorescence using a plate reader (Biotek Synergy2) at 480 nm (excitation) and 528 nm (emission) wavelengths with 20 nm bandwidth. The maximum quench/fluorescence was obtained with 1% Triton-X100 or Tween-20. Each condition was tested in experimental triplicate and the resulting averaged fluorescence was obtained.

### Cell surface biotinylation

Cell surface biotinylation, pulldown and subsequent immunoblotting were performed as described previously (32). Briefly, HEK293 cells were plated onto a 6-well plate and transfected using JetPrime (Polyplus) with 2.5 μg of FLAG-tagged pIE2-Panx1 constructs. Two days post-transfection, cells were harvested and washed in PBS. Surface membrane proteins were biotin-labeled by resuspending cells with 0.5 mg/ml sulfo-NHS-SS-biotin (Thermo Scientific) for 40 minutes at 4 °C. The reaction was quenched by washing cells twice with PBS supplemented with 50 mM NH4Cl, followed by a final wash with 1 mL PBS. Cells were lysed in RIPA buffer (150 mM NaCl, 3 mM MgCl2, 1% NP-40, 0.5% deoxycholate (Anatrace), 0.1% SDS, 20 mM HEPES pH to 7.4 with NaOH) supplemented with 1x protease inhibitor cocktail (Thermo Scientific) and rotated for 30 minutes. The lysate was clarified by centrifugation at 21,000 x g for 15 minutes and the supernatant was recovered. Streptactin sepharose high-performance resin (GE Healthcare) was added to the lysates and rotated for 2 hours and 30 minutes. Samples were washed 6 times and the biotinylated proteins were eluted by incubating resin with 1.5 x SDS sample buffer supplemented with 75 mM DTT for 30 minutes at 55 °C with intermittent vortexing. Anti-FLAG (1:2000; clone M2), or anti-actin monoclonal antibodies (1:2000; line AC-40), were used to detect the target proteins by western blot.

### TEV protease purification

His-6x TEV protease (S219V) was recombinantly expressed and purified using BL21 (DE3) E.coli cells. Briefly, transformed BL21 (DE3) cells were grown to an OD of 2.0-4.0 in Terrific Broth at 37 °C with 250 rpm shaking and expression was induced with 1mM IPTG. The temperature was reduced to 30°C and expression proceeded for 4 hours. Cells were collected via centrifugation and lysed with a combination of freeze-thaw, 0.5 mg/ml lysozyme, and sonication in lysis buffer (2x PBS, 10% glycerol, 0.5mM TCEP, 0.5mM phenylmethylsulfonyl fluoride). Lysate was incubated in DNase (0.1 mg/ml) and MgCl2 (10mM) with stirring for 10 minutes. Lysate was cleared by high-speed centrifugation at 31,000 xg for 45 minutes. TEV protein was bound to TALON resin for 1 hour with stirring, then washed with 20 column volumes of lysis buffer supplemented with 5mM imidazole. Protein was eluted with lysis buffer and 200mM imidazole. Solution was dialyzed in 2x PBS, 20% glycerol and spiked with 0.5mM TCEP before storing.

### frPanx1 purification

P2 baculoviruses of both frPanx1-ΔC (frPanx1 ending at amino acid 357 with an internal Strep II tag) and frPanx1-ΔC+CAD (frPanx1 ending at amino acid 382 with an internal Strep II tag) were amplified in Sf9 cells and 5 mL was used to infect HighFive cells at 3.0x10^6^ cells/mL. Forty-eight hours after infection, cells were harvested by centrifugation and broken using nitrogen cavitation

(4635 cell disruption vessel; Parr Instruments). Cell membranes were collected by high-speed centrifugation at 185,000 xg for 1 hour. Membranes were homogenized in PBS supplemented with 15% glycerol and a protease inhibitor cocktail (2.0 μg/mL leupeptin, 8.0 μg/mL aprotinin, 2.0 μg/mL pepstatin, and 0.5mM phenylmethylsulfonyl fluoride) using a Dounce Homogenizer operated at 100 rpm followed by solubilization in PBS, 15% glycerol, and 1% C12E8 for 1 hour with stirring. The lysate was cleared via high-speed centrifugation (185,000 xg for 40 minutes) and bound to StrepTactin XT resin (Cytiva; cat. 29401326). The washed and concentrated eluate (in 0.027% C12E8, 20mM Tris-HCl pH 8.0, 150mM NaCl, 50mM Biotin) were subject to size exclusion chromatography on a Superose 6 10/300 GL column (Cytiva; cat. 17517201). Panx1 fractions were retrieved for nanodisc reconstitution. Briefly, a mixture of POPC:POPG:POPE:Cholesterol (Avanti) in reconstitution buffer (50mM HEPES pH 7.4, 150mM NaCl) was sonicated with 0.001% C12E8 and mixed with MSP2N2 at a molecular ratio of 1:68 protein to lipid. The scaffold protein and lipids were allowed to associate for 1 hour on ice. Panx1 was added to the mixture at a final ratio of 1:5:337 Panx1 to MSP to lipid. The reconstitution mixture was incubated on ice for 30 minutes, then diluted below the CMC of C12E8 with reconstitution buffer and added to 0.5 g methanol-activated Biobeads SM-2 (Bio-Rad; 1523920). The mixture was rotated end over end for 18 hours and then subject to room temperature size exclusion chromatography on a Superose 6 10/300 GL column. Fractions were collected and concentrated to 1.5 mg/ml for cryoEM analysis.

### CryoEM Sample Preparation and Data Acquisition

Samples were spotted on glow discharged, Quantifoil 1.2/1.3 holey carbon gold grids with 300 mesh (Electron Microscopy Services; Q3100AR1.3) and plunge frozen using a Mark IV vitrobot with a blot time of 4 seconds and force of 7. Humidity was 100% and the chamber was 15 degrees. Grids were initially screened on a Talos Arctica operated at 200 keV and detected using a K3 direct electron detector equipped with a Gatan Imaging Filter. Grids with suitable ice thickness were then collected at Brookhaven National Lab Laboratory for Biomolecular Structure on a Titan Krios operated at 300 keV and detected using a K3 direct electron detector with a Gatan Imaging Filter set to 15 eV. Automated data acquisition was carried out using EPU (Thermo scientific) a nominal magnification of 81,000x with a superresolution pixel size of 0.535Å.

### CryoEM data processing and model building

Both frPanx1-ΔC and frPanx1-ΔC+CAD datasets were imported into RELION 4.0 (Zivanov et al. 2022) and motion corrected using MotionCorr2 (44), accessed through SBgrid. Motion corrected micrographs were moved to cryoSPARC 4.0 (45) for CTF estimation, where any micrographs with a CTF estimation >5 were purged. Particles were picked using template picker, where the template was generated using frPanx1 (PDB: 6VD7). Inspect particle picks was deployed to reduce the number of non-protein picks. Accepted particles were extracted at a box size of 416 and binned 2x. Particles were 2D classified to remove any junk particles. *Ab initio* maps were generated and used for subsequent rounds of heterorefinement. When the maps stopped improving, the particle stack was moved to RELION for 3D classification without alignments to purge particles that did not contribute to high resolution. A series of 3D classifications with tau fudge parameters from T=4-64 followed by 3D refinement imposing C7 symmetry were performed to select homogeneous particles. The final stack of particles was CTF refined, particle polished using Bayesian polishing, and sharpened with post-processing in RELION. Maps were sharpened with a specified b-factor that did not over-sharpen the N-terminal density but allowed sufficient resolution for model building. To better visualize the loop connecting the NTD and the first transmembrane helix, the protomers were isolated and classified as follows: Panx1 maps were symmetry expanded using the relion_particle_symmetry_expand command. One protomer from this C7 expanded map was carefully extracted with the particle subtraction job using a mask generated from a model of a single pannexin protomer. The mask was expanded by 2 pixels for frPanx1-ΔC and no pixels for frPanx1-ΔC+CAD and was given a 5 pixel and 3 pixel soft edge, respectively. Subtracted particles were classified with 3D classification without alignments. Class distributions were analyzed to determine the populations of conformations within each subunit. The existing frog Panx1 structure (PDB:6VDF)(27) was fit into the cryo-EM density maps for model building in Coot (46). Models were iteratively refined using Phenix Real Space Refine (47) until the refinement parameters stopped improving. Deposited maps and models can be accessed at PDB: 9BZ7 and EMDB: EMD-45055 for frPanx1-ΔC, and PDB: 9BZ8 and EMDB: EMD-45056 for frPanx1-ΔC+CAD.

### Continuum electrostatics calculations

The PDB structures for the NTD flipped down (frPanx1-ΔC) and NTD flipped up (frPanx1-ΔC- CAD) were prepared using the PDB Reader & Manipulator tool in CHARMM-GUI (48). Each construct was modeled as partial segments (frPanx1-ΔC: 11-87, 101-158, 195-357 and frPanx1- ΔC-CAD: 2-91, 101-158, 196-376) with NTER and CTER patching at the respective termini, and default protonation states on all ionizable residues. Both structures were oriented so that the permeation pathway was aligned onto the Z axis, and with residue W104 and W127 marking the ends of the transmembrane helices that align with the two membrane/water interfaces, positioned around Z ≈ -17 Å and Z ≈ 17 Å respectively. Continuum electrostatics calculations were carried out on these structures using the Poisson-Boltzmann solver PBEQ in CHARMM 44b2, using the parameters listed in Table S2 and atomic radii from previous studies by Nina et al. (49).

As described previously (50, 51), the electrostatic contribution to an ion’s interaction inside the pore can be calculated by solving the Poisson-Boltzmann equation and electrostatic potential for three situations: the ion+channel complex, the ion alone and the channel alone. The difference in the free energies is the interaction energy:

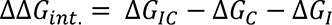

This can also be rewritten in an equivalent form, which collects the contributions in terms of the ion energy in the reaction field, *A*, from the ion charge on the protein and membrane dielectric boundary, the protein static field, B, and the reaction field on the ion dielectric boundary, C:

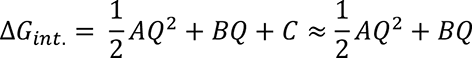

When the channel radius is large, the *C* contribution is negligible. Therefore, the interaction free energy requires solving the Poisson-Boltzmann equation in three situations. The reaction field, *A*, is calculated by solving the potential field for an ion inside the channel, with protein charges turned off, and then subtracting the field created by the ion alone in the high dielectric environment. The static field B is obtained by solving for the field over space without the ion. A summary of these calculations is depicted in Fig. S6.

For the ATP structures, ATP was docked into the cytosolic domain of frPanx1_DC_updated2.pdb using the Autodock Vina extension of the Samson software suite. After adding hydrogen atoms, the ATP structure was energy minimized prior to docking, using the energy minimization function in Samson. All ATP bonds were set to be rotatable, and frPanx1_DC was kept rigid (no flexible sidechains). The search exhaustiveness parameter was set to 32 to predict the top 10 most favorable binding sites. The default value was used for all other search parameters. PBEQ calculations were carried out as described above, with 0-4 ATP left docked in the cytosolic vestibule. Hybrid structures where varying numbers of NTDs docked inside the pore were built by manually building the selected subunits in CHARMM after alignment of both structures such that W104 and W127 flank the membrane interfaces.

## ACKNOWLEDGMENTS

We thank the Kawate lab members for discussion. This work was supported by National Institutes of Health grant R01GM114379 and Cornell Margaret and Richard Riney Canine Health Center Research Grants Program. The Robertson lab is supported by National of Institutes Health grants R01GM120260 and R03NS133680. This work relied on data collected using an instrument supported by the NIH through award S10OD030470. This work made use of the Cornell Center for Materials Research Shared Facilities which are supported through the NSF MRSEC program (DMR-1719875). The Laboratory for BioMolecular Structure (LBMS) is supported by the DOE Office of Biological and Environmental Research (KP1607011).

## AUTHOR CONTRIBUTIONS

E.H., J.L.R., and T.K. designed research; E.H., J.J.E., and J.L.R. performed research; All authors contributed to analyze the data and write, review, and edit the manuscript.

## DATA SHARING PLAN

Structure coordinates and EM density maps have been deposited in the Protein Data Bank (PDB) and the Electron Microscopy Data Bank (EMDB) with accession codes 9BZ7 and EMD- 45055 for frPanx1-ΔC, and 9BZ8 and EMD-45056 for frPanx1-ΔC+CAD. All other data are included in the manuscript or in Supporting Information.

---

**Fig. S1.**
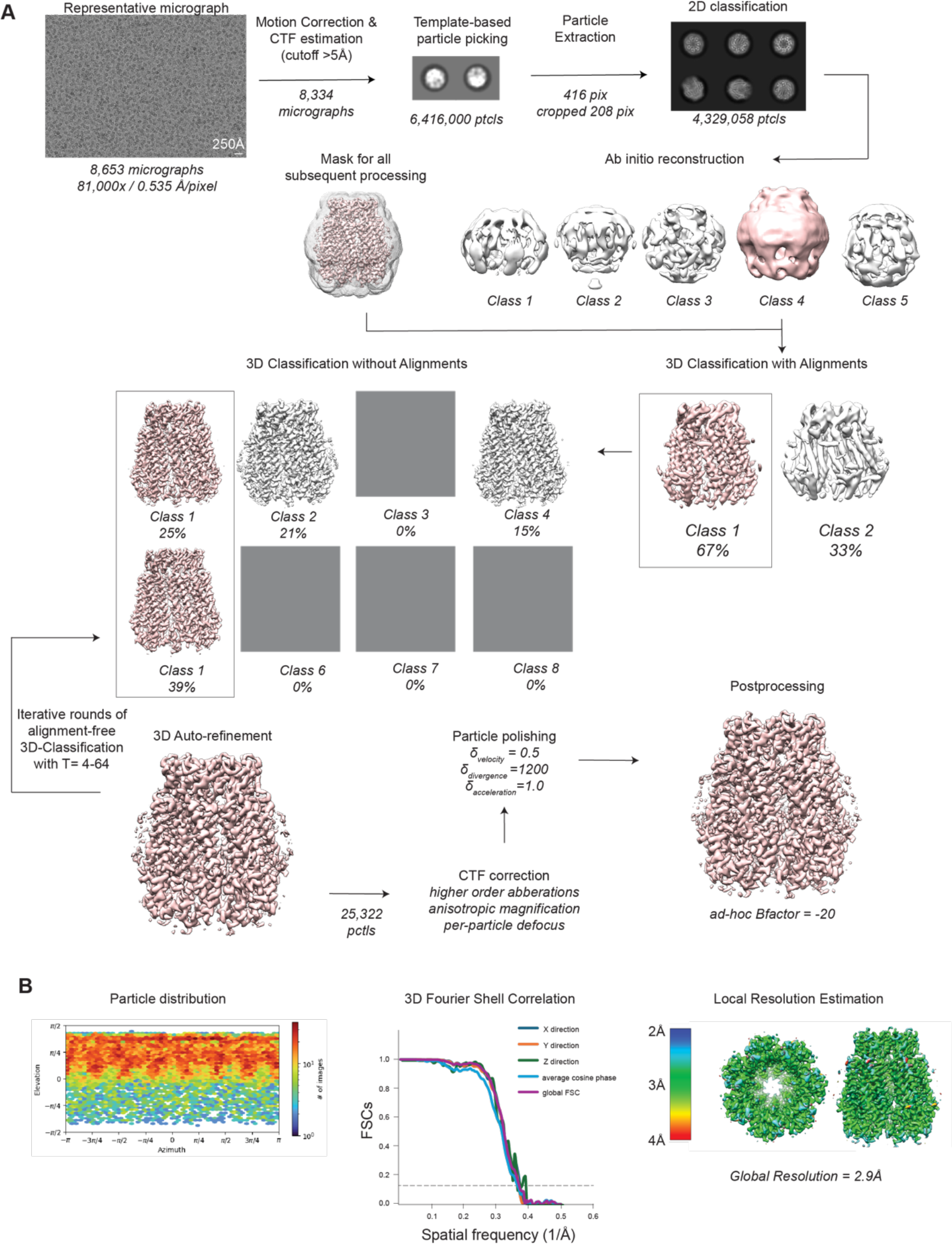
(A) Cryo-EM data processing workflow for frPanx1-ΔC. Initial micrograph processing was carried out in Relion 4.0 (52), then moved to cryoSPARC 4.0 (45) for 2D classification. Particles were moved back to Relion for 3D classification and final map refinement. Model fitting was performed with Coot (46) and model refinement was done with Phenix. Tau-fudge parameters executed in sequence to isolate highest resolution particles are T=4, T=8, T=16, T=32, and T=64 with a 3D-autorefinement job in between each. (B) Refinement statistics for frPanx1-ΔC maps. Particle distribution and local resolution (cutoff 0.143) maps from cryoSPARC. 3DFSC charts were generated via the remote 3DFSC processing server hosted at the New York Structural Biology Center (53).

**Fig. S2.**
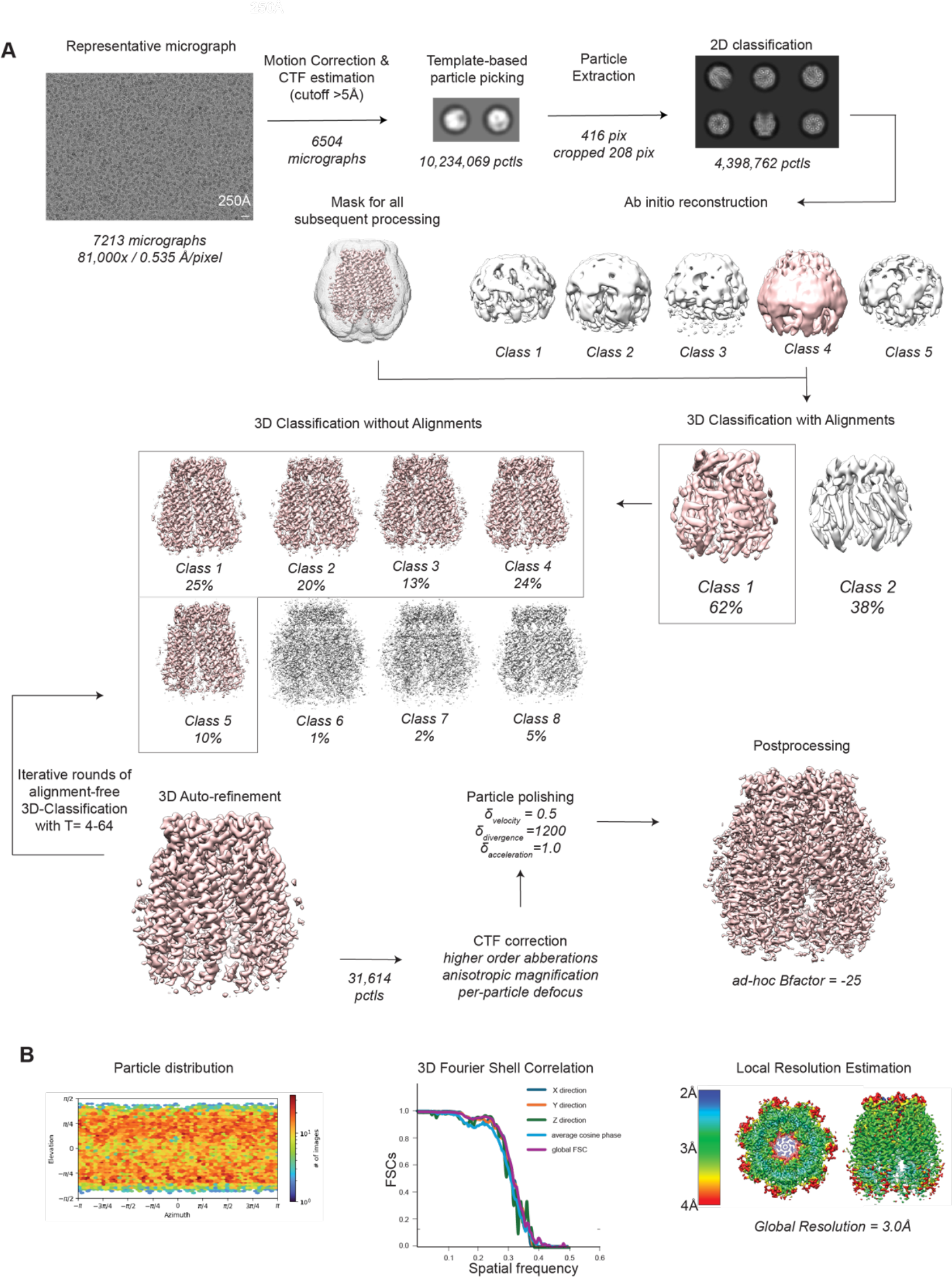
(A) Cryo-EM data processing workflow for frPanx1-ΔC+CAD. Initial micrograph processing was carried out in Relion 4.0 (52), then moved to cryoSPARC 4.0 (45) for 2D classification. Particles were moved back to Relion for 3D classification and final map refinement. Model fitting was performed with Coot (46) and model refinement was done with Phenix. Tau-fudge parameters executed in sequence to isolate highest resolution particles are T=4, T=8, T=16, T=32, and T=64 with a 3D-autorefinement job in between each. (B) Refinement statistics for frPanx1-ΔC+CAD maps. Particle distribution and local resolution (cutoff 0.143) maps from cryoSPARC. 3DFSC charts were generated via the remote 3DFSC processing server hosted at the New York Structural Biology Center (53).

**Fig. S3:**
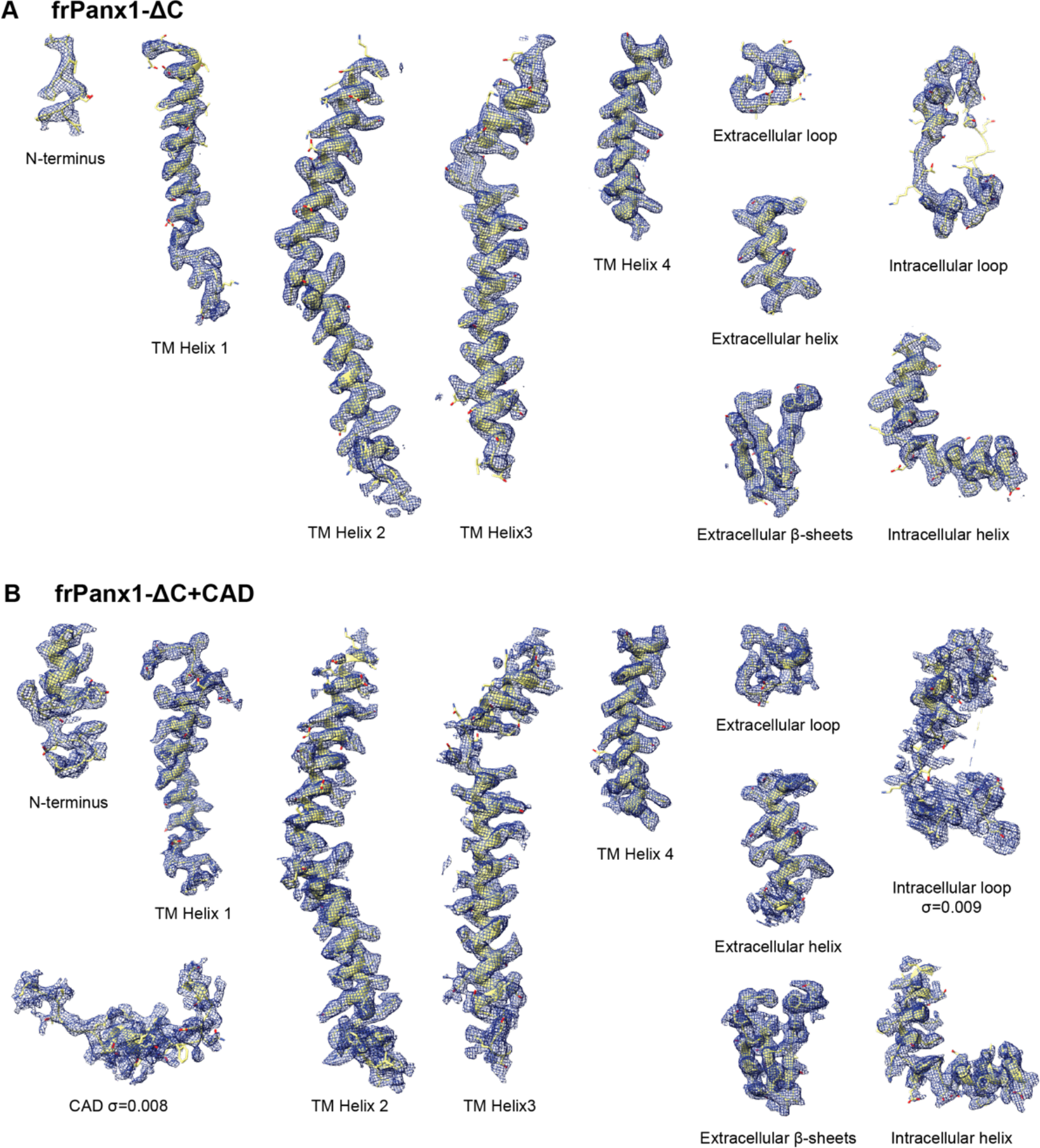
**Cryo-EM density at different parts of frPanx1**. The models are fit into the cryo-EM maps of frPanx1-ΔC (A) and frPanx1-ΔC+CAD (B). Thresholds for maps are 0.01 and 0.013, respectively, unless otherwise noted.

**Fig. S4:**
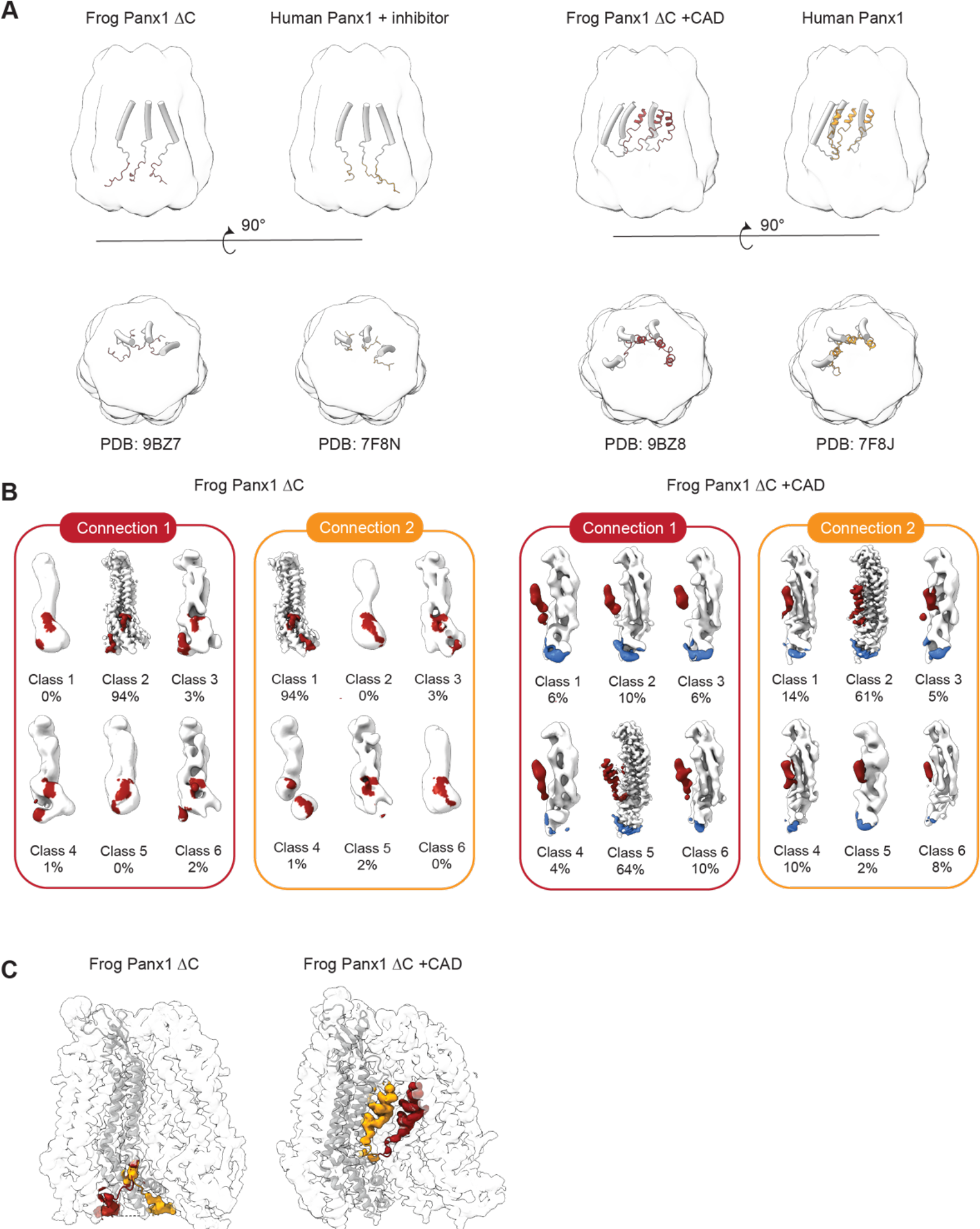
**Structural comparison of the Panx1 structures.** (A) Structures of human and frog Panx1 orthologs have consistent architecture in closed and open or primed conformations with a heptameric assembly and the narrowest constriction located at W74. Structures obtained from this study are shown in red and previously reported structures from Kuzuya et al (30) are shown in blue. Top view is from the extracellular side perpendicular to the plasma membrane and side views are taken parallel to the membrane. (B) Cryo-EM data processing results of single protomers from 3D classification. Maps were generated for each N-terminal connection and the same batch of particles were subject to particle subtraction and 3D classification to better visualize the density connecting the N-terminus to the first TM helix. (C) Comparison of the possible N-terminal positions in both constructs. Density highlighted in red was used for modeling the structures from this paper, and densities highlighted in orange is the reported connection by Kuzuya et al.

**Fig. S5:**
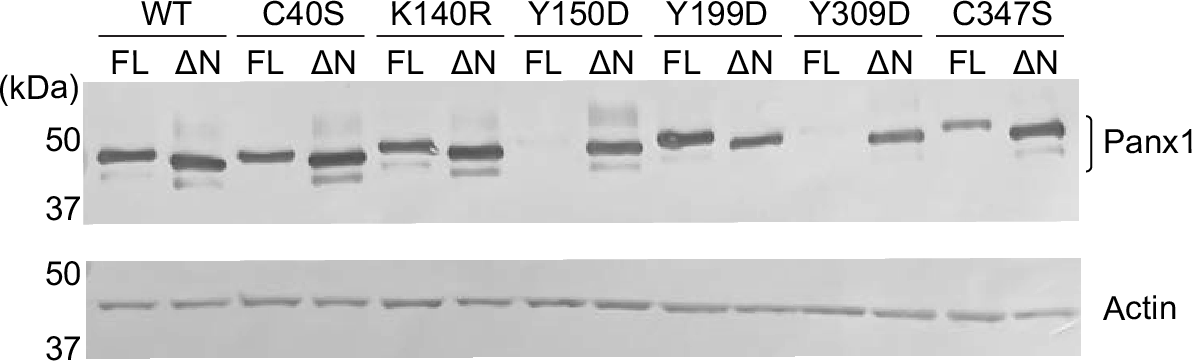
**Surface expression levels of PTM mutants.** Surface-biotinylated Panx1 PTM mutants were pulled down and detected by western blot using anti FLAG antibody. Anti-actin antibody was used to control the loading amount. The target proteins were detected by colorimetric method using alkaline phosphatase substrates.

**Fig. S6:**
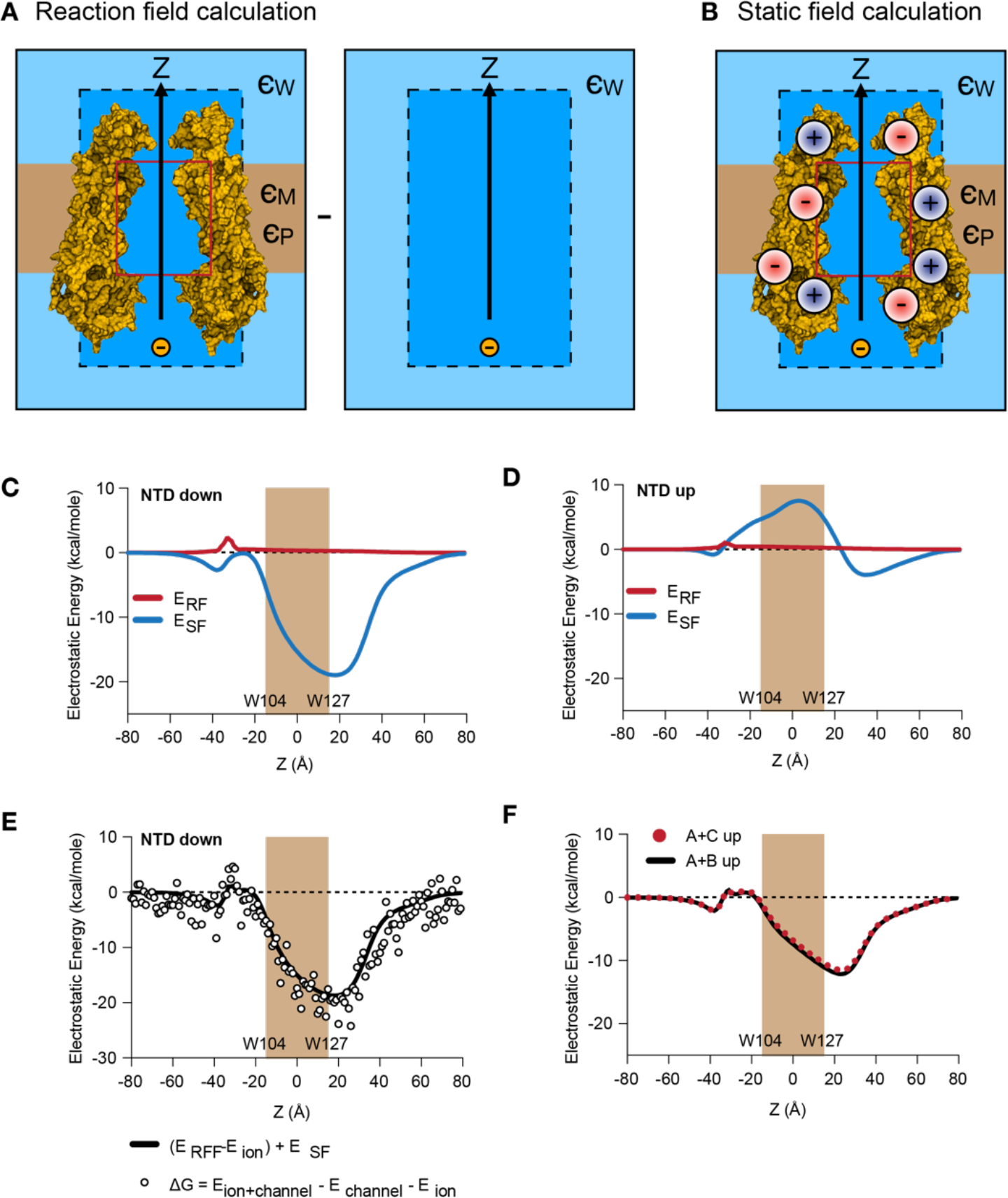
**Continuum electrostatics calculations.** (A) In a continuum Poisson-Boltzmann calculation, the water/protein/membrane system is defined along a 3D grid and solved using the PBEQ module in CHARMM. The water, protein and membrane dielectrics are represented as *ɛW*, *ɛP*, and *ɛM* respectively. In addition, there is a cylinder representation (red box) that allows for the dielectric environment inside the pore to be set to ɛW. The dashed line defines an additional box, so that ionic screening is considered only outside of the pore. The solid arrow indicates the axis of ion permeation, defined along Z. The reaction field contribution to the electrostatic interaction free energy is calculated by solving for the ion in the field of the protein/membrane dielectric environment with all charges turned off (*ERFF*) and subtracting the ion alone in the box *(Eion)*, where *A = ERFF-Eion*. (B) The protein static field contribution is calculated by solving the field in 3D space due to the protein charges in the dielectric environment. (C) The energy due to the reaction field is calculated *as ERF = ½ AQ^2^* and static field *ESF = BQ^2^*. *ERF* and *ESF* for the NTD down structure of frPanx1 and (D) NTD up structure that includes the CAD. (E) A comparison of the two approaches to calculating the interaction energy of the ion inside of the pore, *ERF+ESF* (line) and *ΔGint. = Eion+channel – Echannel - Eion* (points), showing overlap. The noise in the latter calculation arises from variability in the solution when moving the ion along *Z* in the *Eion+channel* calculation. (F) Comparison of *ERF+ESF* in the hybrid models where A+B vs. A+C subunits were made with the NTD in the up position, showing similar electrostatic profiles.

**Fig. S7:**
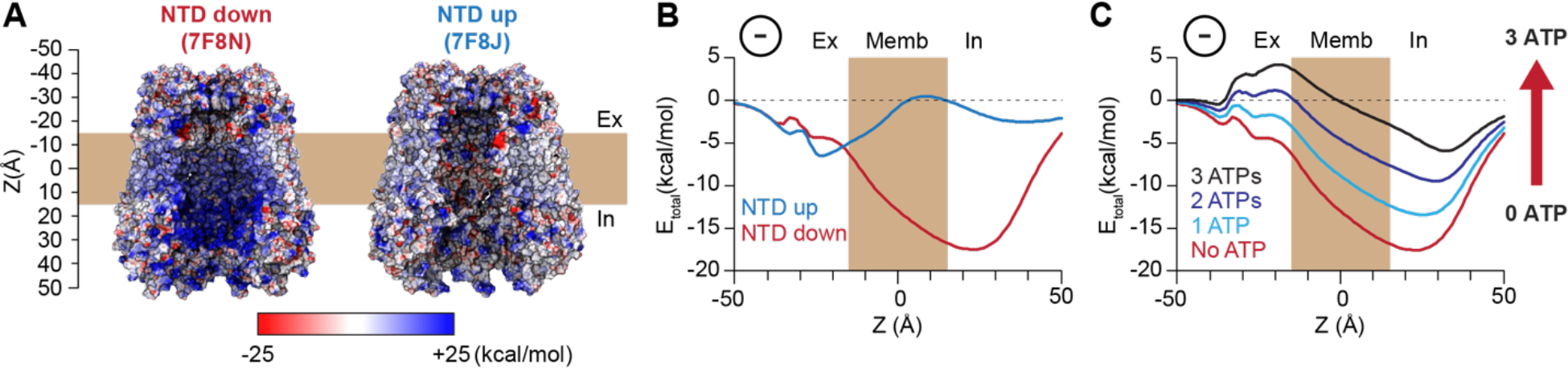
**Electrostatic energy landscape of the human Panx1 channel.** (A) Electrostatic surface potential of a coronal section of the frPanx1-ΔC (left) and frPanx1-ΔC+CAD (right) ion permeation pathway. Electrostatic surface potential was calculated using CHARMm-PBEQ (1) and presented in the range between -25 kcal/mol (red: acidic) and +25 kcal/mol (blue: basic). (B) Electrostatic contribution for an anion binding free energy (Etotal) along the permeation pathway (z=0 in the middle of predicted transmembrane) for the 7F8N (red) and 7F8J (blue), and hybrid models harboring 0-7 NTDs (C) Electrostatic free energy calculation of 7F8N (red) in the presence of 0-3 ATP molecules in the deep energy well. Calculations are carried out using CHARMM-PBEQ module to solve the Poisson-Boltzmann solution of the system.

**Table S1.** Statistics for reconstructions and model building

| Codes | frPanx1-ΔC<br>PDB:<br>EMDB: | frPanx1-ΔC+ CAD<br>PDB:<br>EMDB: |
| --- | --- | --- |
| **Data collection and processing** |  |  |
| Microscope | Titan Krios | Titan Krios |
| Camera | K3 BioQuantum | K3 BioQuantum |
| Magnification | 81,000x | 81,000x |
| Voltage (kV) | 300 | 300 |
| Defocus Range | -1.0 to -2.0 | -1.0 to -2.0 |
| Exposure time | 3.81 | 3.83 |
| Total dose | 50 | 50 |
| Number of frames | 50 | 50 |
| Pixel size | 1.07 (0.535 in super-resolution) | 1.07 (0.535 in super-resolution) |
| Micrographs (no.) | 8334 | 6504 |
| Symmetry imposed | C7 | C7 |
| Initial particles (no.) | 6,416,000 | 10,234,069 |
| Final particles (no.) | 25,322 | 31,614 |
| Map resolution (Å) | 2.9 | 3.0 |
| FSC threshold | 0.143 | 0.143 |
| **Refinement** |  |  |
| Initial model (PDB code) | 6VD7 | 6VD7 |
| Model resolution (Å) | 3.1 | 3.0 |
| FSC threshold | 0.5 | 0.5 |
| Model composition |  |  |
| Non-hydrogen atoms | 16,961 | 18,277 |
| Protein residues | 2,086 | 2,254 |
| Ligands | 0 | 0 |
| B factors (Å<sup>2</sup>) | -20 | -25 |
| R.m.s deviation |  |  |
| Bond length (Å) | 0.002 | 0.002 |
| Bond angle (°) | 0.544 | 0.511 |
| **Validation** |  |  |
| Favored (%) | 95.89 | 97.77 |
| Allowed (%) | 4.11 | 2.23 |
| Disallowed (%) | 0 | 0 |
| Rotamer outliers (%) | 1.84 | 2.73 |
| MolProbity score | 1.95 | 1.79 |
| Clash score | 8.74 | 7.34 |

**Table S2.** Parameters for PBEQ calculations. All other parameters are set to default values.

| Variable | Value | Physical meaning |
| --- | --- | --- |
| (NCELX,NCELY,NCELZ) | frPanx1-ΔC: (289, 287, 397) | Number of grid points |
| DCEL1 | 1.0 Å | Coarse grid dimension |
| DCEL2 | 0.5 Å | Focusing grid dimension |
| (XBCEN,YBCEN,ZBCEN) | (0 Å, 0 Å, 0 Å) | Center of box |
| EPSP | 2 | Protein dielectric constant |
| EPSW | 80 | Water dielectric constant |
| CONC | 0.15 M | Ionic concentration where allowed |
| WATR | 1.4 Å | Solvent probe radius |
| IONR | 0.0 Å | Ion exclusion radius |
| TEMP | 300 K | Temperature |
| ZMEMB | 0 Å | Center of membrane |
| TMEMB | 30 Å | Membrane thickness |
| EPSM | 2 | Membrane dielectric constant |
| RCYLN | 15 Å | Radius of cylinder to form aqueous pore across membrane |
| HCYLN | 30 Å | Height of cylinder across membrane |
| EPSC | 80 | Cylinder dielectric |
| (XCYLN,YCYLN,ZCYLN) | (0 Å, 0 Å, 0 Å) | Center of cylinder |
| (LXMIN,LYMIN,LZMIN) | (-25 Å, -25 Å, -55 Å) | Minimum box dimensions |
| (LXMAX, LYMAX,LZMAX) | (25 Å, 25 Å, 65 Å) | Maximum box dimensions |
| EPSB | 80 | Box dielectric (except protein) |

